# Among retroviral integrases prototype foamy virus integrase displays unique biochemical activities

**DOI:** 10.1101/2021.11.17.469013

**Authors:** Anthony J. Rabe, Yow Yong Tan, Ross C. Larue, Kristine E. Yoder

## Abstract

Integrase enzymes of different retroviruses assemble as functional complexes with varying multimers of the protein. Retroviral integrases require a divalent metal cation to perform one-step transesterification catalysis. Tetrameric prototype foamy virus (PFV) intasomes assembled from purified integrase and viral DNA oligonucleotides were characterized for their activity in the presence of different cations. While most retroviral integrases are inactive in calcium, PFV intasomes appear to be uniquely capable of catalysis in calcium. The PFV intasomes also contrast other retroviral integrases by displaying an inverse correlation of activity with increasing manganese beginning at relatively low concentrations. The intasomes were found to be significantly more active in the presence of chloride co-ions compared to acetate. While HIV-1 integrase appears to commit to a target DNA within 20 seconds, PFV intasomes do not commit to target DNA during their reaction lifetime. Together these data highlight the unique biochemical activities of PFV integrase compared to other retroviral integrases.

## Introduction

Retroviruses require reverse transcription and integration to complete the viral life cycle [1]. Reverse transcriptase copies the viral genomic RNA to a linear double stranded DNA (cDNA). Integrase (IN) acts on the viral cDNA with two necessary catalytic activities: terminal cleavage and strand transfer. Terminal cleavage, also called 3’ end processing, is the removal of two nucleotides from the 3’ ends of nascent reverse transcripts yielding recessed 3’ hydroxyls. Strand transfer is the covalent joining of these 3’ hydroxyls to the host DNA. The 3’ ends are joined across one major groove of host DNA with 4-6 base pairs of intervening host sequence. The product of IN activities in vivo is the integrated provirus flanked by 4-6 base gaps of host sequence and 5’ dinucleotide flaps of viral sequence. The integration intermediate is likely repaired by host DNA repair enzymes [2]. Retroviral IN may also catalyze intramolecular strand transfer, termed autointegration. This phenomenon effectively ends a cellular infection and may also appear during biochemical assays. Half site integration (HSI) products where a single viral DNA is joined to a target DNA has also been observed in vitro. It is unclear if these aberrant integration products occur under normal conditions in vivo, although observed altered integration sites suggest they may occur with mutant IN or in the presence of suboptimal concentrations of clinically relevant IN strand transfer inhibitors [3,4].

Retroviral INs catalyze 3’ processing and strand transfer by single step transesterification chemistry [5-7]. Each step may be independently assayed with DNA oligomers mimicking blunt viral cDNA ends (vDNA) to test 3’ processing or preprocessed vDNA to test strand transfer. These enzymes do not require an energetic co-factor such as ATP [8]. INs have a conserved DDE motif that coordinates 2 divalent metal ions. IN and similar enzymes, including transposases, have been characterized for their use of metal cations. Magnesium (Mg) and calcium (Ca) are the most abundant of these cations in a cell, but Ca is more highly regulated within cellular compartments [9]. Similar to Mg, Ca is often able to bind the catalytic site of enzymes, but is unable to support catalysis [9,10]. The reason for enzymatic inactivity or altered activity with Ca is mysterious and several hypotheses have been postulated including variation in ion size or coordination number [9-16]. Manganese (Mn) is less abundant than Mg but enzymes are often able to employ this cation for catalysis [10,15,17]. Although enzymes may be active with Mn, the activity may be altered from that observed with Mg, such as less stringent sequence specificity [11,12,16,18]

Human immunodeficiency virus (HIV-1) IN was characterized for its ability to assemble with vDNA, perform 3’ processing, or catalyze strand transfer to a target DNA in the presence of Mn, Mg, Ca, cobalt (Co), or no cation [19]. These results revealed that HIV-1 IN may assemble an integration complex in the presence of Ca, but could not complete 3’ processing or strand transfer. HIV-1 IN assembled with vDNA in the presence of Ca could perform enzymatic activities when Mg or Mn were added. This observation has been fundamental for the recent advances in assembly and purification of stable retroviral intasomes, including HIV-1 and mouse mammary tumor virus (MMTV) [20,21]. Purified IN combined with vDNA oligomers in the presence of Ca can form intasomes, the catalytically active multimeric IN complex. The intasomes are stable for purification by size exclusion chromatography and do not perform autointegration. This strategy allows for the precise control of catalytic activity with later addition of Mg to purified intasomes. Interestingly, prototype foamy virus (PFV) intasomes do not require the presence of a divalent cation for assembly [22,23].

Previous studies using monomeric protein suggested HIV-1 IN and avian sarcoma virus (ASV) IN displayed greater catalytic activity in the presence of Mn compared to Mg [24]. Similarly, PFV IN was shown to have greater strand transfer activity yielding more HSI and CI products in the presence of Mn but only with a blunt vDNA, suggesting that Mn stimulated 3’ processing activity [25]. However, HSI and CI with a preprocessed vDNA were equal in the presence of Mn or Mg. These data indicate that PFV IN 3’ processing is more active in the presence of Mn, but strand transfer is not. As seen with IN from other retroviruses, PFV IN was inactive in the presence of Ca and a blunt vDNA indicating it was unable to perform 3’ processing [25]. PFV IN displayed an unexpected ability to perform strand transfer generating HIS products with preprocessed vDNA in the presence of Ca. This suggested PFV IN can perform assembly and strand transfer in the presence of Ca, but not 3’ processing. It was unclear whether intasomes would display the similar results in the presence of these divalent metal ions, particularly since PFV intasomes typically generate less HSI products compared to PFV IN.

Retroviral INs have been extensively studied with biochemical assays of recombinant protein and vDNA oligomers, but fewer studies have employed intasomes. PFV intasome assembly and purification was the first to be described [22]. These intasomes were visualized by crystallography and shown to include a tetramer of IN [22]. PFV intasomes readily perform integration to a plasmid target DNA in vitro [22]. Here we extend the biochemical characterization of PFV intasomes. PFV intasome product saturation occurred in 2 min. The kinetics are not affected by the molar ratio of IN to target DNA, the addition of a small molecule that prevents intasome aggregation, or the presence of a second target DNA. PFV IN appears unique among retroviral INs in the capacity to utilize calcium as a divalent cation for strand transfer. Finally, HIV-1 IN commits very quickly to a target DNA, but PFV IN and intasomes do not. These enzymatic properties distinguish PFV IN from other retroviral INs.

## Materials and Methods

### Expression and purification of PFV IN

PFV IN was purified as previously described [26,27]. Briefly, hexahistidine tagged PFV IN was induced in *E. coli* strain BL21(DE3) pLysS with 250 μM IPTG at 25°C for 4 h. Soluble cellular lysate was fractionated by nickel affinity chromatography. Fractions with PFV IN were treated with HRV 3C protease to remove the hexahistidine tag. PFV IN was further purified by heparin affinity chromatography. Fractions containing highly concentrated PFV IN were combined, dialysed against 50 mM Tris HCl, pH 7.5, 500 mM NaCl, 5 mM DTT, and 10% glycerol, aliquoted, snap frozen with liquid nitrogen, and stored at -80°C.

### Assembly and purification of PFV intasomes

Intasomes were assembled as previously described [23]. Briefly, 50 mM Bis-tris propane, pH 7.5, 500 mM NaCl, 120 μM PFV IN, and 50 μM vDNA were combined in a total volume of 150 μL and dialyzed overnight at 18°C against 20 mM Bis-tris propane, pH 7.5, 200 mM NaCl, 2 mM DTT, and 25 μM ZnCl_2_. The intasome aggregates were solubilized by increasing the concentration of NaCl from 200 mM to 320 mM and incubating on ice. The intasomes were purified by size exclusion chromatography using a Superose 12 10/300 (GE Healthcare) equilibrated with 20 mM Bis-tris propane, pH 7.5, 320 mM NaCl, and 10% glycerol. Fractions containing intasomes were aliquoted, snap frozen with liquid nitrogen, and stored at -80° C. PFV intasomes appear to retain activity for one year at -80°C.

### Integration Assays

Standard integration reactions were performed in 30 mM Bis-tris propane, pH 7.5, 110 mM NaCl, 5 mM MgSO_4_, 4 μM ZnCl_2_, 10 mM DTT, indicated concentration of PFV intasomes and 1.8 nM supercoiled plasmid target DNA in a final volume of 15 μL. Reactions were incubated for 5 min at 37° C, stopped with the addition of 0.5% SDS, 0.5 mg/mL proteinase K, 25 mM EDTA (pH 8.0), and incubated for 1 h at 37°C. Integration products were resolved with 1.25% agarose gel electrophoresis. Gels were stained with ethidium bromide and imaged for ethidium bromide and Cy5 fluorescence (Sapphire Biomolecular Imager, Azure Biosystems). Cy5 fluorescence was quantified using AzureSpot gel analysis software (Azure Biosystems). Where indicated 5 mM PCA was included in reactions. Acetate buffer included 25 mM Tris-HCl, pH 7.4, 125 mM NaOAc, 5 mM MgOAc, 10 μM ZnCl_2_, 1 mM DTT. In assays comparing buffers, 20 nM PFV intasome was added to reaction buffer and incubated on ice for indicated times before addition of 1.8 nM 3 kb supercoiled pGEMT plasmid. At the addition of target DNA, reactions were incubated at 37°C for 5 min and analyzed as standard integration reactions. P values were determined by two tail paired t test. Error bars indicate standard deviation between at least three independent experiments performed with at least two independent intasome preparations.

## Results

### PFV intasome requirements for divalent cations

Retroviral intasomes may be assembled with a preprocessed vDNA and purified recombinant PFV IN [22]. The vDNA may be labelled with a fluorophore, such as Cy5, to identify integration products by fluorescence imaging [23]. The assembled complexes are purified by size exclusion chromatography. To assay integration activity the PFV intasomes are diluted in a buffer containing a divalent cation and target DNA. Addition of supercoiled plasmid as the target DNA allows facile visualization and quantitation of the integration products. Concerted integration (CI_1_) of both vDNAs to a circular plasmid yields a linear product with vDNA at the ends (Figure 1). Additional CI (CI_2_) to the linear product results in fragments shorter than the linear product. Half site integration (HSI) is the joining of a single vDNA to the plasmid and results in a fluorescently tagged plasmid with the mobility of a relaxed circle. Autointegration (AI) products result from integration of one vDNA to another vDNA and have slightly slower mobility compared to unreacted vDNA. These reaction products are resolved by agarose gel electrophoresis. The agarose gel is imaged for ethidium bromide and Cy5 fluorescence and quantified. Integration products are distinguished by their mobilities and presence of the fluorophore.

**Figure 1.**
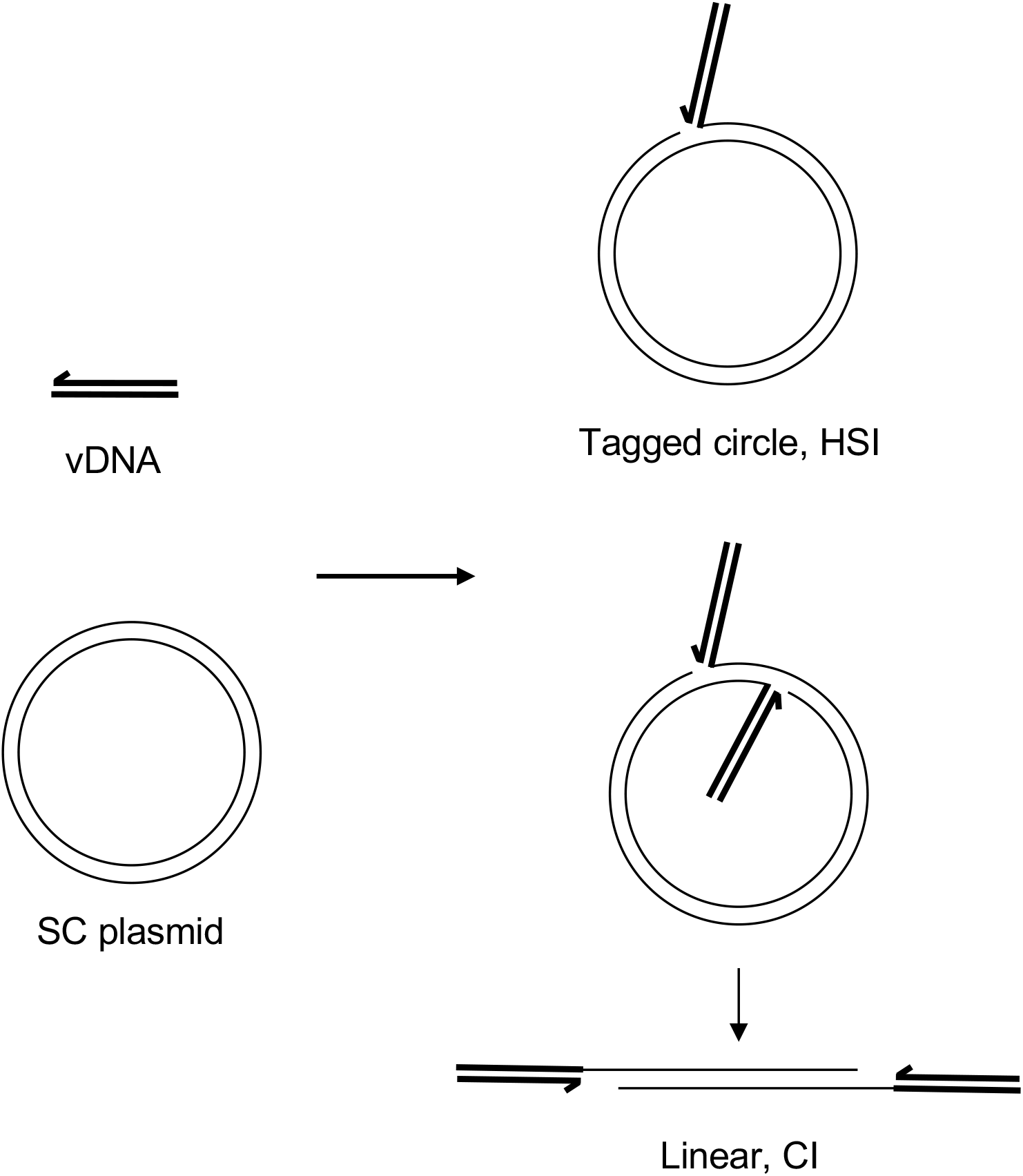
Integration assay reaction products. DNA oligomers mimicking the viral cDNA ends (vDNA, heavy black lines) are assembled with PFV IN to form intasomes. The intasomes are added to a supercoiled plasmid (SC plasmid, light black lines) and incubated at 37°C. The major products of integration are concerted integration (CI) where the two vDNAs of the intasome are covalently joined to the target SC plasmid. This results in a linear product with the vDNAs at each end. Additional CI to the linear product will result in shorter fragments. A minor product of intasome integration is half site integration (HSI) where only one vDNA is joined to the target DNA resulting in a tagged circle.

PFV intasomes with preprocessed vDNA were assembled without a divalent metal ion. The intasomes were purified by size exclusion chromatography in the continued absence of divalent metal ion indicating that the complexes are stable without cations. The use of a preprocessed vDNA allows analysis of strand transfer activity. PFV integration assays are commonly performed in the presence of magnesium sulfate [25,28]. PFV intasome integration to a supercoiled plasmid was performed with a titration of magnesium sulfate (Figure 2A). In the absence of cation PFV intasomes are unable to perform strand transfer. Maximal concerted integration activity was observed in the presence of 5 mM magnesium sulfate. Similar results were seen with a titration of magnesium chloride (Figure 2B). There was no statistically significant difference between PFV integration activity in the presence of magnesium sulfate or magnesium chloride (p > 0.05 at all equivalent concentrations).

**Figure 2.**
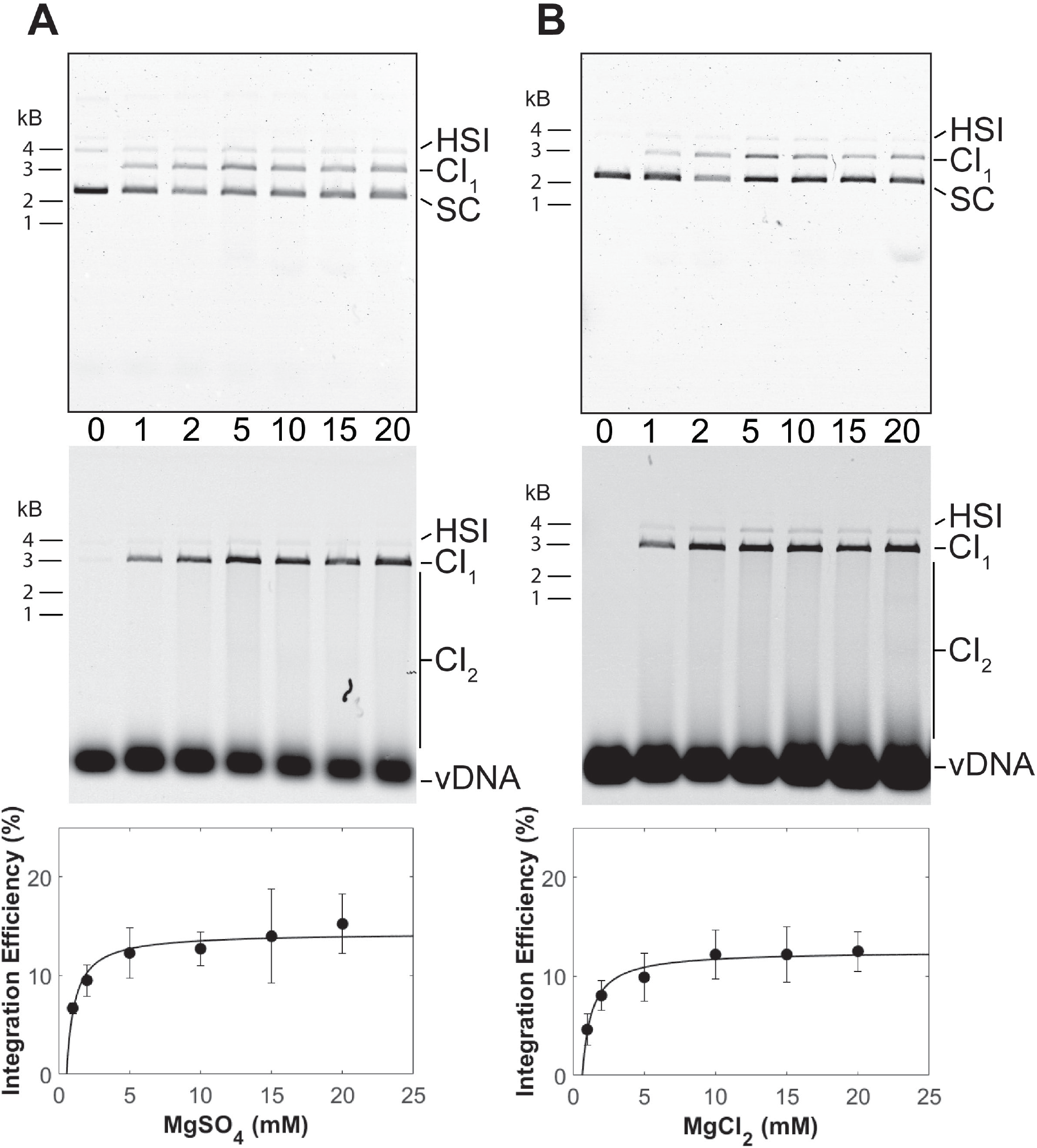
PFV intasome integration in the presence of Mg. Integration activity was assayed with a titration of (**A**) MgSO_4_ or (**B**) MgCl_2_. Reaction products were resolved by agarose gel electrophoresis. The gel was imaged for ethidium bromide (top) and Cy5 (center) fluorescence. Half site integration (HSI), the first concerted integration product (CI_1_), and subsequent concerted integration products (CI_2_) are shown. Unreacted supercoiled plasmid target DNA (SC) is visible on in the ethidium bromide image. Unreacted vDNA is only visible in the Cy5 image. The total Cy5 fluorescence in each lane was quantified and each integration product calculated as the fraction of the total fluorescence (bottom). The average of three independent experiments with at least two independent intasomes preparations is shown. Error bars indicate the standard deviation.

PFV intasomes were also assayed for integration activity in manganese chloride and calcium chloride. Intasomes were most active in the presence of 1 mM manganese chloride (Figure 3). In contrast to magnesium, increasing concentration of manganese chloride led to decreased integration activity. Previous studies of HIV-1 IN also revealed an inhibition of strand transfer, but only at 64 mM manganese chloride [29]. PFV intasomes were also assayed with a titration of calcium chloride (Figure 4). Many retroviral integrases are not active in the presence of calcium. However, PFV intasomes were active in the presence of calcium chloride and activity increased with increasing concentrations of calcium chloride. In the presence of this divalent metal cation integration products were mostly HSI. Employing a higher concentration of PFV intasomes revealed the formation of CI products in the presence of Ca. These data highlight the unique characteristics of PFV IN divalent metal requirements compared to other retroviral INs.

**Figure 3.**
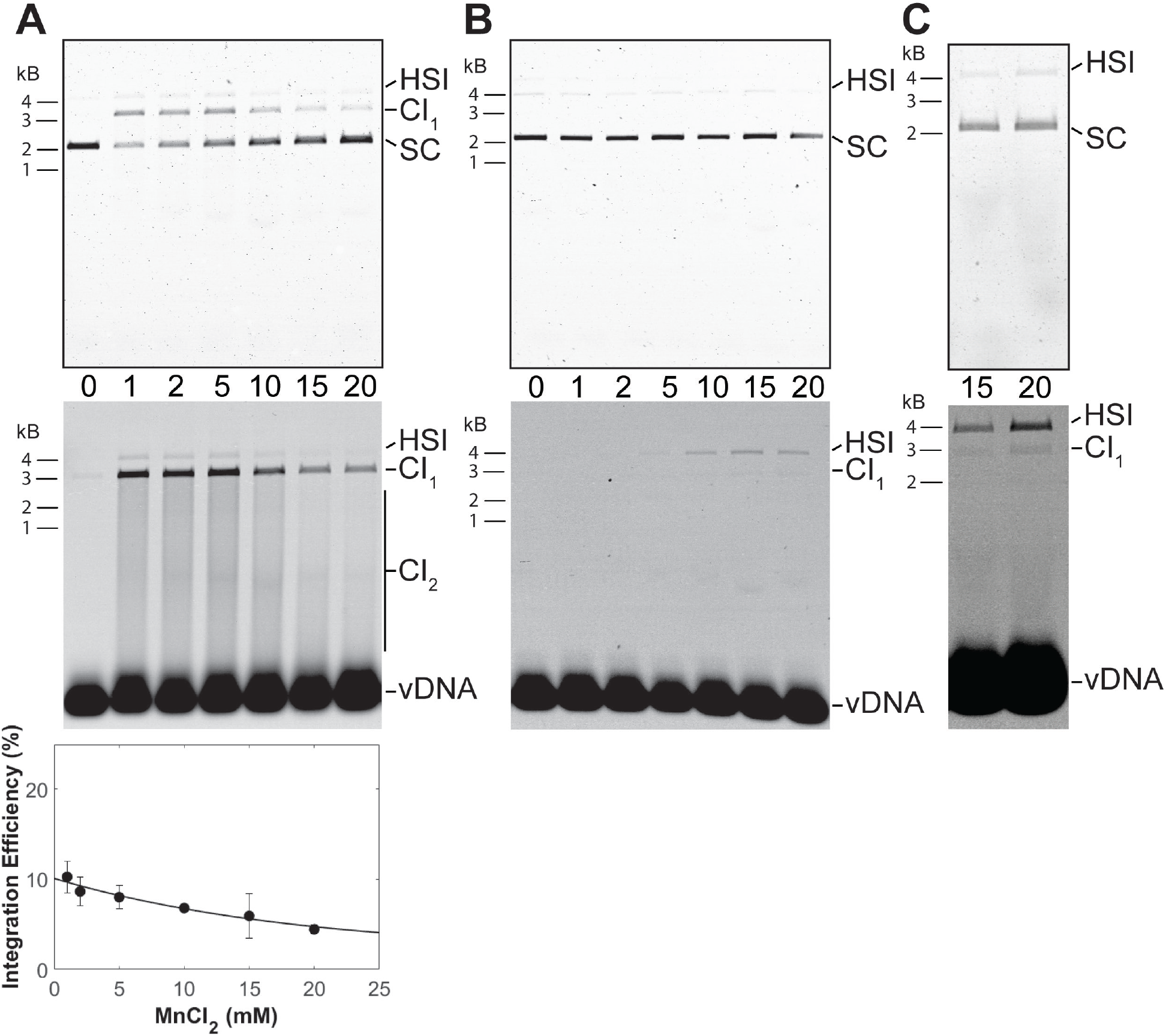
PFV intasome integration in the presence of MnCl_2_ or CaCl_2_. (**A**) Integration activity was assayed with a titration of MnCl_2_ and 10 nM PFV intasomes. Reaction products were resolved by agarose gel electrophoresis. The gel was imaged for ethidium bromide (top) and Cy5 (center) fluorescence. Half site integration (HSI), the first concerted integration product (CI_1_), and subsequent concerted integration products (CI_2_) are shown. Unreacted supercoiled plasmid target DNA (SC) is visible on in the ethidium bromide image. Unreacted vDNA is only visible in the Cy5 image. The total Cy5 fluorescence in each lane was quantified and each integration product calculated as the fraction of the total fluorescence (bottom). The average of three independent experiments with at least two independent intasomes preparations is shown. Error bars indicate the standard deviation. (**B**) 10 nM PFV intasomes were assayed for integration with a titration of CaCl_2_. Reaction products were resolved by agarose gel electrophoresis. The gel was imaged for ethidium bromide (top) and Cy5 (center) fluorescence. HSI and CI_1_ are shown. Unreacted SC target DNA is visible on in the ethidium bromide image. Unreacted vDNA is only visible in the Cy5 image. The reaction products were < 1% of the fluorescent signal in each lane and could not be accurately quantified. (**C**) A higher concentration of PFV intasomes, 45 nM, was assayed to confirm the presence of CI products in the presence of Ca.

**Figure 4.**
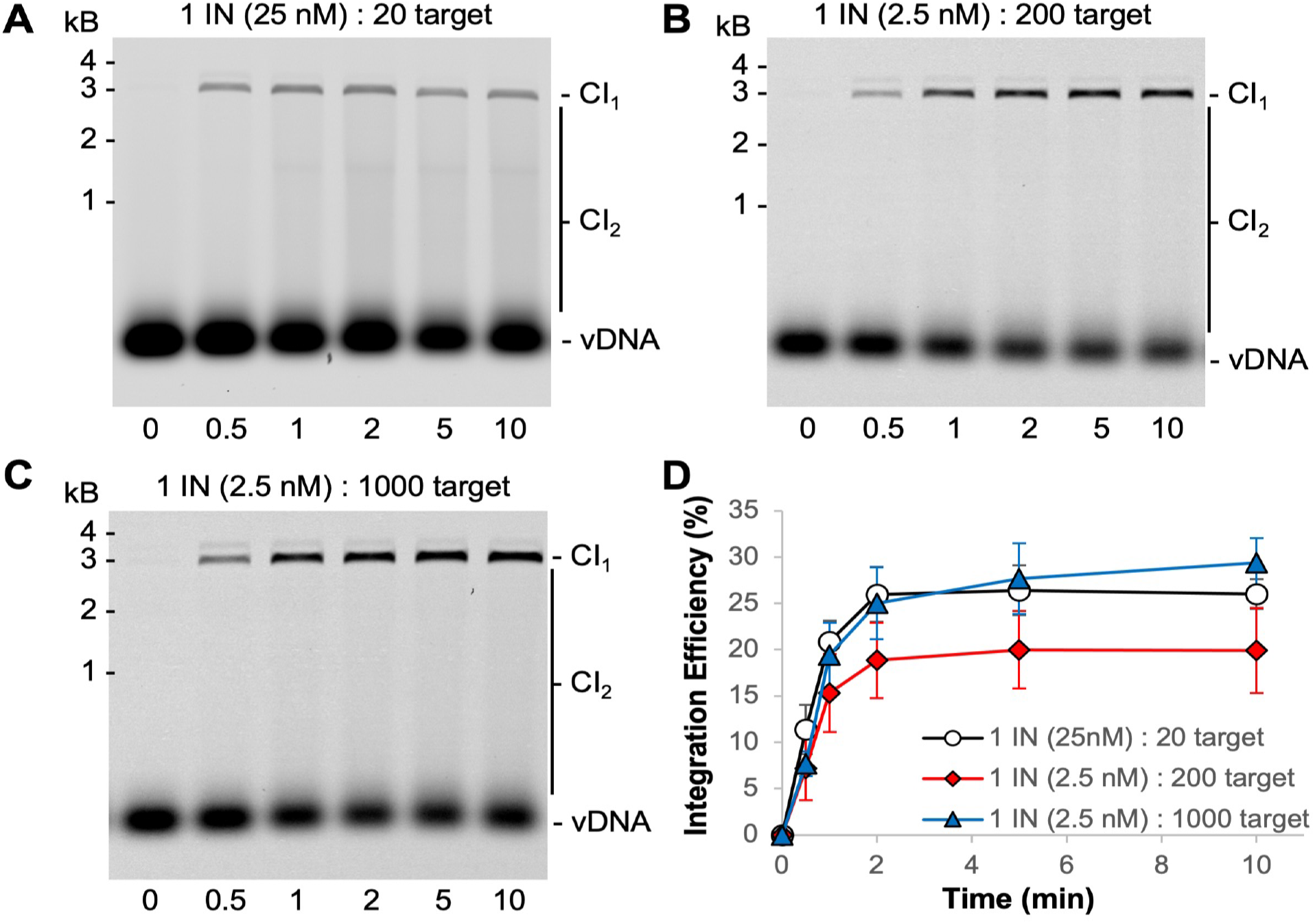
Increasing the molar ratio of target to PFV intasomes. The footprint of the PFV intasome is ∼10 bp suggesting that 3000 bp plasmid target offers roughly 300 possible integration sites. (**A**) Typical reaction conditions with 25 nM PFV intasomes is a 20 fold molar excess of target sites to intasomes. (**B**) Reducing the intasome concentration to 2.5 nM increases the molar excess of target sites to 200 fold. (**C**) Increasing the target DNA concentration allowed for a 1000 fold molar excess of target sites to 2.5 nM intasomes. Cy5 images of agarose gels show the accumulation of CI_1_ and CI_2_ products over time. (**D**) The total Cy5 fluorescence in each lane was quantified and each integration product calculated as the fraction of the total fluorescence. The average of three independent experiments with at least two independent intasomes preparations is shown. Error bars indicate the standard deviation.

### PFV intasome mediated integration is quick

PFV intasomes were previously shown to complete integration within 5 minutes at 37°C with either a supercoiled plasmid target or nucleosome target [30,31]. However, there was only one earlier time point assayed in those studies. For a more accurate determination of intasome kinetics, we extended the time course with the same buffer conditions and found that integration is complete within 2 minutes at 37°C (Figure 4).

The PFV intasome footprint appears to be ∼10 bp of target DNA [32]. A 3000 bp plasmid target offers 300 possible integration sites. Integration assays were performed with a molar excess of target sites to PFV intasomes (Figure 4) [30,33]. Reactions with 25 nM PFV intasomes are at a 20 fold molar excess of target sites to intasomes. The intasome concentration was reduced to 2.5 nM in order to evaluate a 200 fold molar excess of target sites. The 200 fold molar excess reactions yielded less CI products, particularly CI_2_ products, compared to 20 fold molar excess of target sites. All reactions were complete by 2 min. The amount of target plasmid included in the reactions was increased to obtain a 1000 fold molar excess of target sites to 2.5 nM intasomes. The data with altered molar ratios of target substrate to intasomes consistently show the reaction to be complete at 2 min. Neither the substrates nor the intasomes were completely consumed during these reactions.

The short reaction time of PFV intasome mediated integration makes classic Michaelis-Menten analysis of enzyme kinetics difficult. Previous studies showed that PFV intasomes incubated at 37°C for 5 min before the addition of target lost integration activity [30]. However, addition of the small molecule PCA significantly rescued the activity of the pre-incubated PFV intasomes. This was at least in part attributed to the ability of PCA to prevent aggregation of PFV intasomes at physiologically relevant ionic strength buffer conditions [30]. PCA was added to PFV intasomes to test the ability of this small molecule to lengthen the time that intasomes are active beyond 2 min (Figure 5). Two PFV intasome concentrations were assayed in the presence or absence of PCA. The presence of PCA had no effect on the accumulation of CI products over time and reaction products saturated by 2 min. This data suggests that while PCA prevents the aggregation of PFV intasomes, the completion of integration at 2 min was not due to aggregation of complexes.

**Figure 5.**
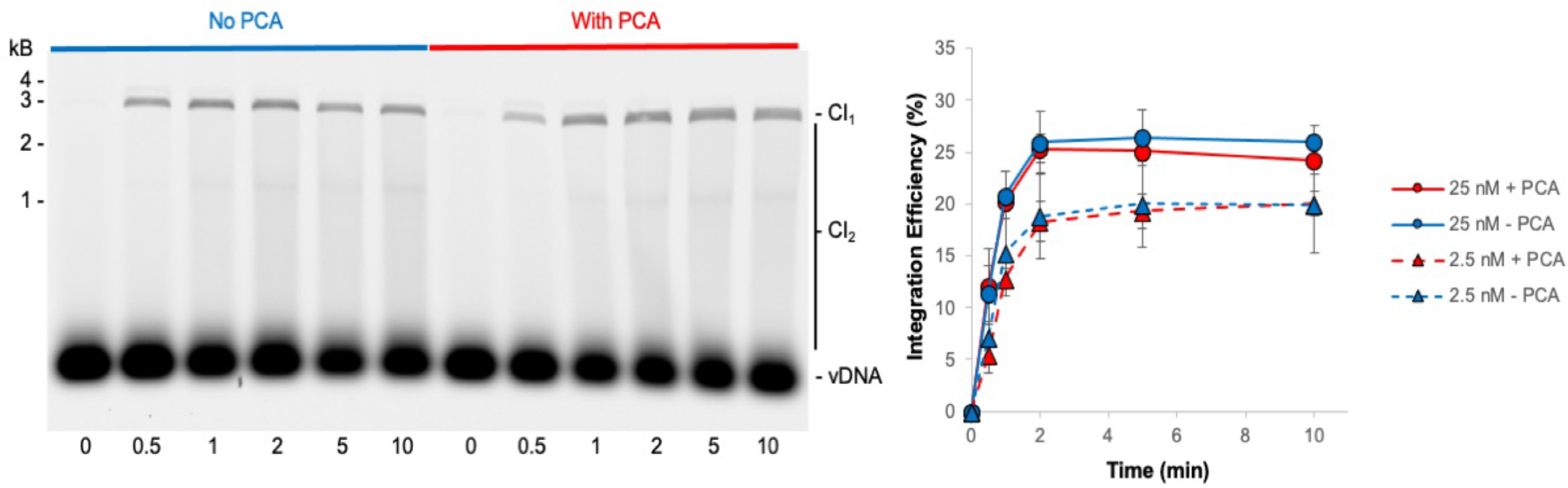
PCA does not alter the kinetics of PFV intasome integration. PFV integration over time was assayed in the absence or presence of 5 mM PCA. As PCA has been shown to reduce aggregation of PFV intasomes, two concentrations of intasomes were assayed: 25 nM and 2.5 nM. Cy5 image of agarose gel with 25 nM PFV intasomes show the accumulation of CI_1_ and CI_2_ products over time. The total Cy5 fluorescence in each lane was quantified and each integration product calculated as the fraction of the total fluorescence. PCA did not affect the kinetics of integration at either concentration. The average of three independent experiments with at least two independent intasomes preparations is shown. Error bars indicate the standard deviation.

### PFV intasomes are less active in the presence of acetate buffer

Studies of PFV intasome activity are routinely performed in the presence of NaCl and MgCl_2_ or MgSO_4_ [25,28,30,33-40]. PFV IN strand transfer showed no difference in the presence of MgSO_4_ or MgCl_2_ indicating no significant effects of sulfite or chloride co-ions (Figure 2) [25]. A recent study employed sodium acetate (NaOAc) and magnesium acetate (MgOAc) during assays of PFV intasome activities [41]. This led to results contradicting previously published results obtained with buffer containing NaCl and MgSO_4_ [33]. While Jones et al were able to measure the time between PFV concerted strand transfer events by magnetic tweezers in the presence of NaCl and MgSO_4_, Vanderlinden et al were only able to observe HSI in the presence of NaOAc and MgOAc. To address the difference in experimental outcomes, PFV intasome activity was directly compared in the presence of a buffer with NaOAc and MgOAc to a buffer containing NaCl and MgSO_4_.

PFV intasome integration activity was measured over time in two different buffer conditions (Figure 6A). PFV intasomes were active in both buffers generating readily visualized CI products. However, the intasomes displayed more CI in buffer with NaCl and MgSO_4_. The relative increase of CI product accumulation in the presence of NaCl and MgSO_4_ is apparent at 0.5 min. In addition, accumulation of CI products plateaus in the presence of NaCl and MgSO_4_ at 2 min incubation. However, CI is complete at 1 min in the acetate buffer. These data indicate that PFV intasomes are less active in acetate buffers.

**Figure 6.**
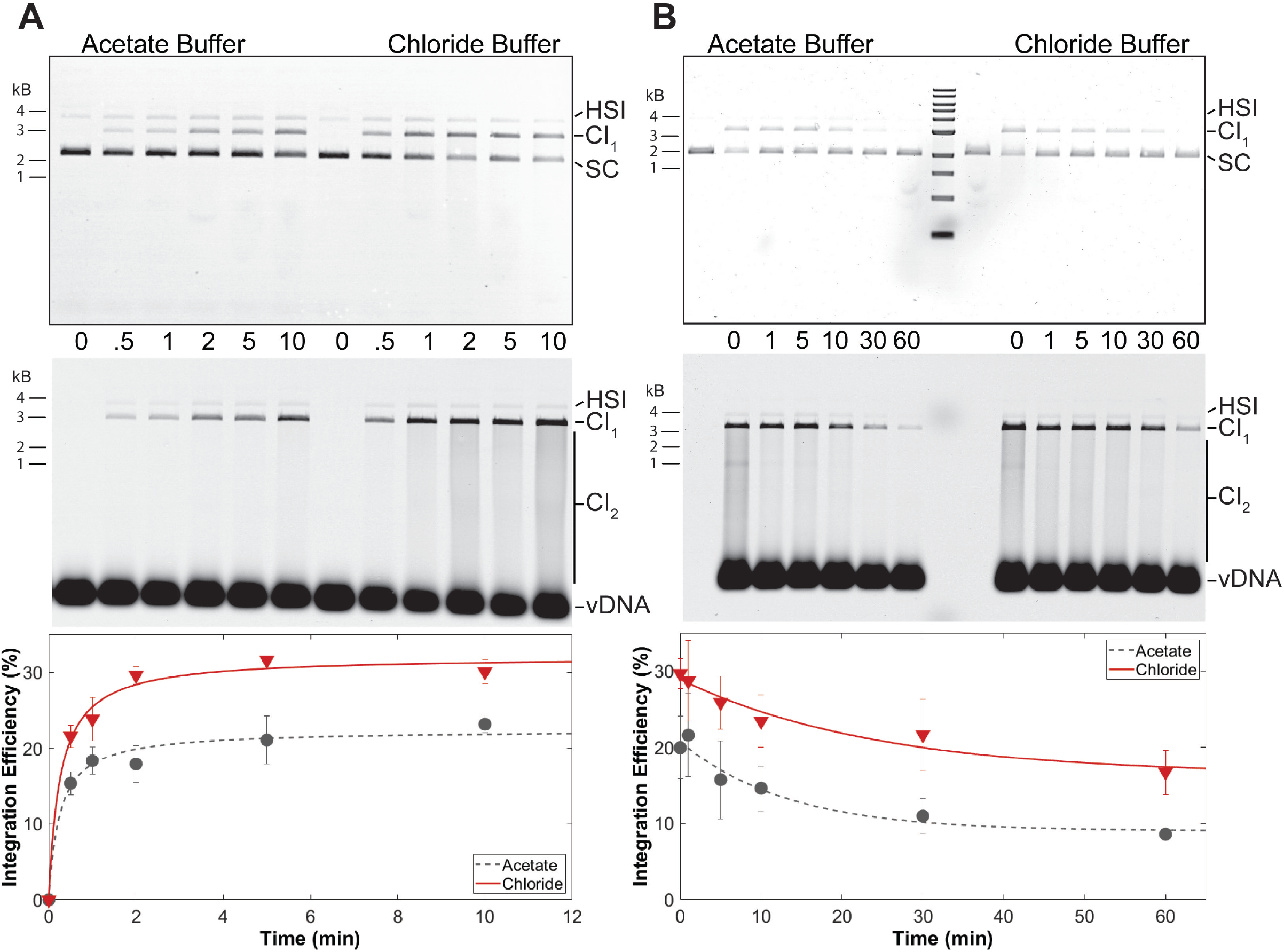
PFV intasomes are more active in chloride buffers. (**A**) PFV integration over time was assayed in the presence of standard chloride buffer or an acetate buffer. (**B**) PFV intasomes were diluted in chloride or acetate buffer and kept on ice for variable time before the addition of target DNA and immediate transfer to 37°C. Reaction products were resolved by agarose gel electrophoresis. The gel was imaged for ethidium bromide (top) and Cy5 (center) fluorescence. Half site integration (HSI), the first concerted integration product (CI_1_), and subsequent concerted integration products (CI_2_) are shown. Unreacted supercoiled plasmid target DNA (SC) is visible on in the ethidium bromide image. Unreacted vDNA is only visible in the Cy5 image. The total Cy5 fluorescence in each lane was quantified and each integration product calculated as the fraction of the total fluorescence (bottom). The average of three independent experiments with at least two independent intasomes preparations is shown. Error bars indicate the standard deviation.

The quick reaction kinetics of PFV intasomes suggested that practical considerations may also play a role in integration assays. In the case of single molecule magnetic tweezers assays, PFV intasomes are typically diluted in reaction buffer before loading to a flow cell. In some cases, the diluted intasomes may remain on ice before being exposed to target DNA within the flow cell. The two different reaction buffers were tested for their effects on intasomes incubated on ice (Figure 6B). PFV intasomes were diluted to a working concentration in reaction buffer and incubated on ice for variable time. Following variable incubation time on ice, target DNA was added and the reactions were immediately transferred to 37°C for 5 min. PFV intasome incubation on ice led to reduced activity over time in both buffers. However, the intasomes in acetate buffer lost 50% of their integration activity after 30 min and 55% after 60 min on ice. Intasomes in buffer with NaCl and MgSO_4_ lost 37% of their activity after 30 minutes and 42% after 60 min of incubation on ice. These data indicate that PFV intasomes are more prone to loss of activity in acetate buffer compared to buffer with NaCl and MgSO_4_.

### PFV intasomes do not commit to target DNA

HIV-1 IN was previously shown to quickly commit to a target DNA [42]. In these assays HIV-1 IN and vDNA were added to a plasmid and incubated at 37°C. At variable times a second plasmid of differing size was added to the reaction. HIV-1 IN performed equivalent integration to both plasmids when they were added simultaneously. However, HIV-1 IN appeared to fully commit to the first plasmid within 20 seconds with no integration to a second plasmid. In contrast, similar experiments showed that PFV IN integrated to a second plasmid up to 60 min after the addition of the first plasmid [25]. Both of these experimental approaches employed free IN and vDNA rather than assembled intasomes.

We tested the commitment of PFV intasomes to two plasmid DNA targets, 3 kb and 6 kb (Figure 7). Reactions were performed with 2.5 nM PFV intasomes to reduce the amount of CI_2_ products which would confound quantitation. When the plasmids were added to the reaction simultaneously, integration to the plasmids was equivalent. As seen with a single plasmid target, integration was complete by 2 min. At times shorter than 2 min, PFV intasomes integrated to either plasmid. Over time the fraction of integration to the first plasmid increased and integration to the second plasmid decreased. The integration dynamics were unaffected by whether the smaller or larger plasmid was added first. These results suggest that PFV intasomes do not fully commit to a target DNA early, as seen with free HIV-1 IN.

**Figure 7.**
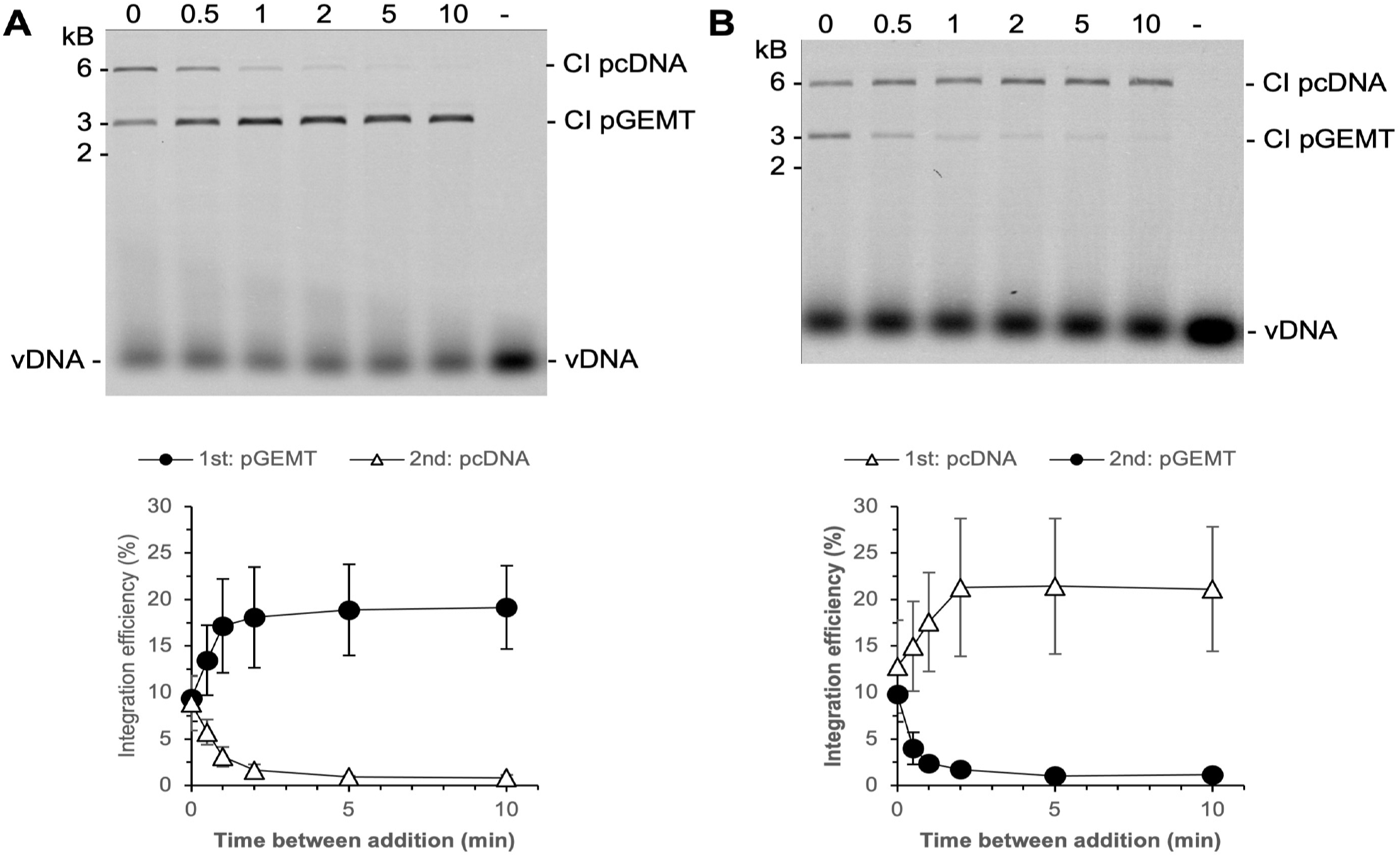
PFV intasomes do not commit early to target DNA. Integration reactions were performed with a 3 kb and a 6 kb plasmid. Reactions were started with one plasmid and a second plasmid added at variable time later. The zero time point represents plasmids added simultaneously to the reaction. (**A**) The 3 kb plasmid was added first and the 6 kb plasmid added second. (**B**) The 6 kb plasmid was added first and the 3 kb plasmid was added second. The total Cy5 fluorescence in each lane was quantified and each integration product calculated as the fraction of the total fluorescence (bottom). The average of three independent experiments with at least two independent intasomes preparations is shown. Error bars indicate the standard deviation.

## Discussion

Many enzymes that utilize two divalent metal cations for catalysis display activity in Mg or Mn, but not Ca. The mechanism of this preference is not clear, but several hypotheses have been proposed. The reaction energy barrier in the presence of Ca appears to be raised for both RNaseH and BamHI [9,14]. Another factor may be the larger atomic radius of Ca compared to Mg [10]. Observations of RNaseH suggested that the architecture of the active site was the same in the presence of Ca or Mg, but changes in ion coordination geometry were observed [9]. This may be a key factor considering the structural similarities of retroviral intasomes at the active site in what has been termed a conserved intasome core (CIC) [43]. Structural studies of PFV intasomes in the presence of Mg or Mn described the octahedral coordination for both binding sites, but Ca was not included in the analysis [44].

It is unclear why PFV IN of all retroviral INs appears uniquely able to utilize Ca in strand transfer catalysis. HIV-1 IN has been the most widely studied for its activity in the presence of various divalent cations. Divalent metals were shown to be required for assembly of the integration complex, 3’ end processing, and strand transfer. HIV-1 IN was shown to efficiently use Mg or Mn for all three activities [19,29]. However, HIV-1 IN could only use Ca for assembly, not 3’ processing or strand transfer [19]. It could use Co for strand transfer, but not assembly or 3’ processing [19]. Similarly, ASV IN performs strand transfer in the presence of Mg or Mn, with better efficiency in the latter [24]. In contrast, PFV intasomes do not require any divalent cation for assembly [22,32]. This enzyme can use Mg or Mn for 3’ processing, but not Ca [25]. However, PFV IN can use Ca for strand transfer (Figure 5) [25].

Multiple transposases have also been characterized for their ability to utilize different divalent cations. Similar to INs, transposases assemble to complexes and perform the same single step transesterification reactions. Phage Mu transposase MuA forms a tetramer and can also perform strand transfer in the presence of Ca [11]. MuA must cleave the Mu DNA ends to form a cleaved donor complex, but is not able to utilize Ca for this activity [11]. This is not true for all transposases as Tn10 transposase has no activity in the presence of Ca [12]. Thus PFV IN activities in the presence of Ca are more similar to MuA transposase than to other retroviral INs.

PFV IN also appears to be distinguishable from HIV-1 IN by a lack of commitment to a target DNA. HIV-1 IN was previously shown to commit to a target DNA within 20 sec [42]. PFV IN or intasomes do not display this commitment [25]. PFV intasomes are able to integrate to a second target DNA at any time while they are active (Figure 7). This could be due to 3D searching of target DNAs or relatively slow binding followed by fast reaction kinetics. These experiments did not distinguish these two models. Regardless of the mechanism, the PFV IN interaction with target DNA is demonstrably different from the observed HIV-1 IN fast commitment to target DNA.

These results highlight the unique biochemical characteristics of PFV IN compared to other retroviral INs. All retroviruses display unique patterns of integration to the genome during infection [45]. These results extend that observation showing that retroviral INs also display unique biochemical characteristics. Further biochemical studies of additional retroviral intasomes will reveal their unique properties.

## Author Contributions

Conceptualization, K.E.Y.; methodology, A.J.R. and Y.Y.T.; formal analysis, A.J.R. and Y.Y.T.; writing—original draft preparation, A.J.R., Y.Y.T., and K.E.Y.; writing—review and editing, A.J.R., Y.Y.T., R.C.L., and K.E.Y.; supervision, R.C.L., and K.E.Y.; funding acquisition, K.E.Y. All authors have read and agreed to the published version of the manuscript.

## Data Availability Statement

All relevant data to this study are presented. Additional data may be provided upon request.

